# Poor codon optimality as a signal to degrade transcripts with frameshifts

**DOI:** 10.1101/345595

**Authors:** Miquel Àngel Schikora-Tamarit, Lucas B. Carey

**Affiliations:** Systems Bioengineering Program, Department of Experimental and Health Sciences, Universitat Pompeu Fabra, Barcelona, Spain.

**Keywords:** nonsense-mediated decay, codon bias, frameshifts, mRNA quality, translation

## Abstract

Living organisms are error-prone. Every second a single human cell produces over 100 transcripts with a substitution, frameshift or splicing error. Multiple mRNA quality control pathways exist to degrade these transcripts. Many of these pathways involve co-translational regulation of mRNA stability, such as nonsense mediated decay (NMD) and reduced stability of transcripts with suboptimal codon usage. Recent work has shown the existence of a genetic link between NMD and codon-usage mediated mRNA decay. Here we present new computational evidence that, because the codons following most frameshift errors are suboptimal, removal of mRNAs with such errors may be mediated by degradation of mRNAs with sub-optimal codons. Thus, most transcripts that contain frameshifts are subject to two modes of degradation.

**Author summary:** Frameshifting errors are common and mRNA quality control pathways, such as nonsense-mediated decay (NMD), exist to degrade these aberrant transcripts. Recent work has shown the existence of a genetic link between NMD and codon-usage mediated mRNA decay. Here we present computational evidence that these pathways are synergic for removing frameshifts.

## Introduction

### Frameshifting errors in gene expression

All biochemical pathways are intrinsically stochastic processes. Transcription, splicing, and translation are especially error prone, with error rates 4-6 orders of magnitude higher than that of DNA polymerase (1–6). Such errors can result in single-amino acid substitutions, as well as truncation of the protein due to nonsense mutations or frameshifting errors. The latter can occur due to insertion and deletion events during transcription, splicing errors, and ribosomal slippage during translation **(Figure 1)**.

**Figure 1:**
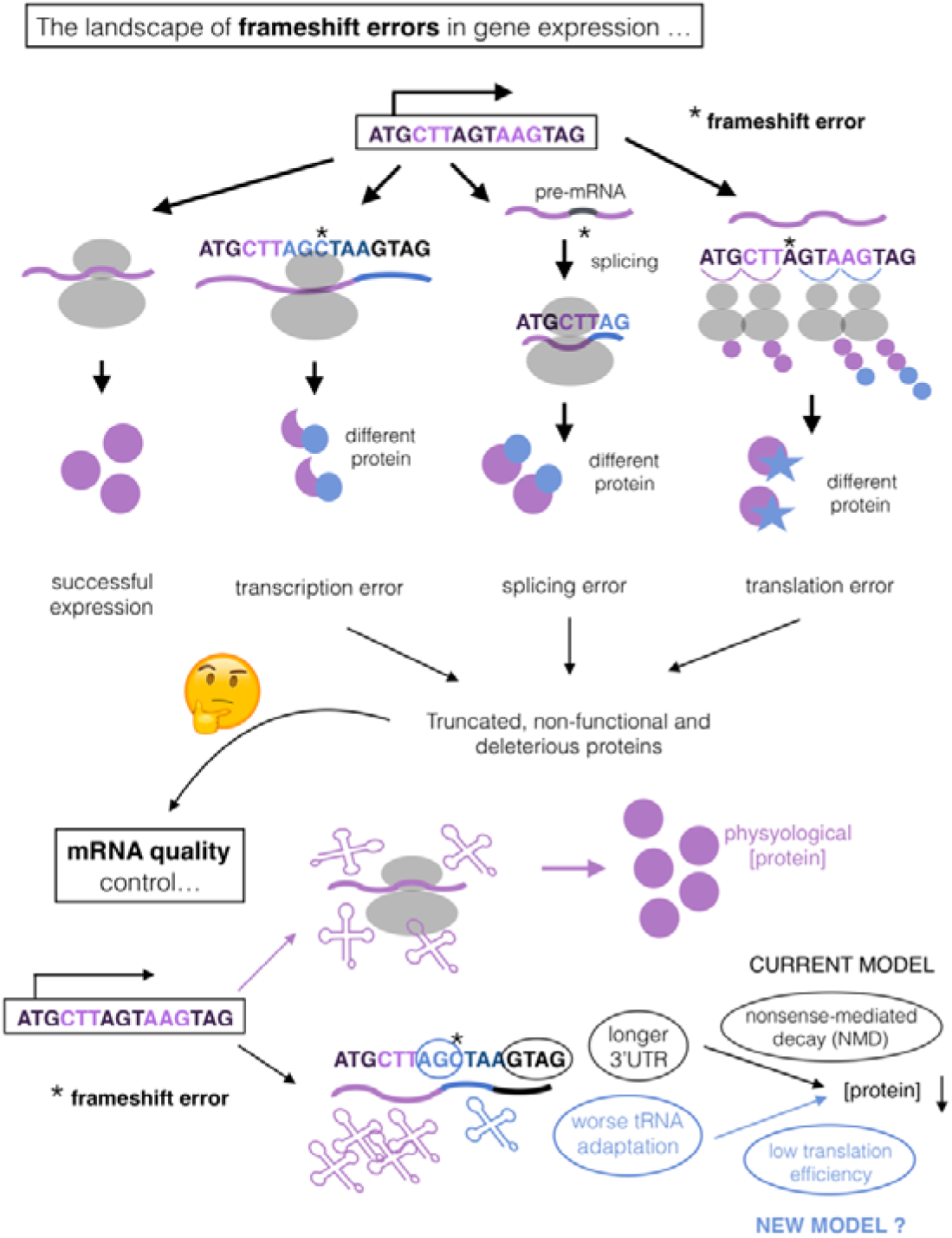
The impact of frameshifting errors in gene expression. Gene expression can result in frameshifting errors (indicated as *) due to transcriptional insertion/deletion epimutations, errors in splicing or ribosomal slippage during translation (top). These processes potentially generate deleterious proteins, which justifies the need of mRNA quality control mechanisms in cells (bottom). In the absence of errors, mRNAs are translated leading to physiological protein levels. The current model indicates that frameshifting errors generate Premature Termination Codons (PTC) that trigger Nonsense-Mediated Decay (NMD) on them, mainly because of the generated long 3’UTR (in yeast). Our hypothesis is that NMD is often nonspecific for errors, so that other quality control mechanisms must exists. We note that another signal of “incorrectness” may appear in transcripts with frameshifts: a stretch of poorly-optimized codons (in blue, indicating worse tRNA adaptation) between the error and the PTC. This should lead to reduced translation efficiency, mRNA decay and lower protein concentrations of the frameshifted transcript.

Frameshifts in protein coding genes are likely to be among the most damaging events, as they result in truncated proteins which may be misfolded or form dominant negative alleles (7,8) **(Figure 1)**. This justifies an evolutionary pressure for cells to contain mRNA surveillance pathways that remove transcripts bearing frameshifts. Suppression of frameshift errors is thought to be one of the major roles of the mRNA quality control machinery (9).

### Nonsense-mediated decay for removing frameshifting errors

In eukaryotes, nonsense-mediated decay (NMD) is a conserved mRNA surveillance pathway that is often assumed to fulfill a frameshift-removing role (10). This follows from the observation that frameshifts generate premature termination codons (PTCs), recognition of which targets the transcript for NMD. However, the quantitative effects of NMD, when measured, are often small (11,12). In addition, a large fraction native transcripts (between 5%-30% depending on the genome) are targeted by NMD (13). In the context of mRNA quality control, these are poor evidence for NMD being an effective quality control pathway.

The mechanism of NMD may be species-specific (10,12) and has even been proposed to be a passive result of the degradation of unprotected transcripts (14). In yeast, NMD is thought to act on long 3’UTRs (15,16), so that transcripts bearing 3’UTRs longer than 250 nucleotides are targeted by NMD **(Figure 1)**. Recent work has shown that this is mostly true and, importantly, the strength of NMD depends linearly on 3’UTR length (11) **(Figure 3B)**. However, native 3’UTR lengths are highly variable, ranging from 0 to 1461 nucleotides (17). Frameshifts in native transcripts with short 3’UTRs are unlikely to result in efficient NMD.

These data suggest that NMD is both inaccurate and inefficient discretizing “correct” vs “incorrect” transcripts. We propose that an efficient quality control pathway should be better able to distinguish and degrade incorrect transcripts.

## Results

### Codon bias and mRNA quality control

Recent work (11) provides an unexpected clue towards understanding mRNA quality control. Two mechanisms of co-translational regulation, NMD and codon bias-dependent mRNA expression (18,19) **(Figure 2A)** are genetically linked; both pathways are regulated by the DEAD-box RNA helicase Dbp2 and by promoter architecture. A quantitative analysis of the impact of these pathways on mRNA levels gives rise to the hypothesis that they may act in a synergistic manner to remove transcripts with frameshifts. In addition to generating a PTC, frameshifts generate a second signal of “wrong transcript”: a run of normally out-of-frame codons between the frameshift and the PTC that are now translated **(Figure 1)**. Below we provide computational support of this hypothesis.

**Figure 2:**
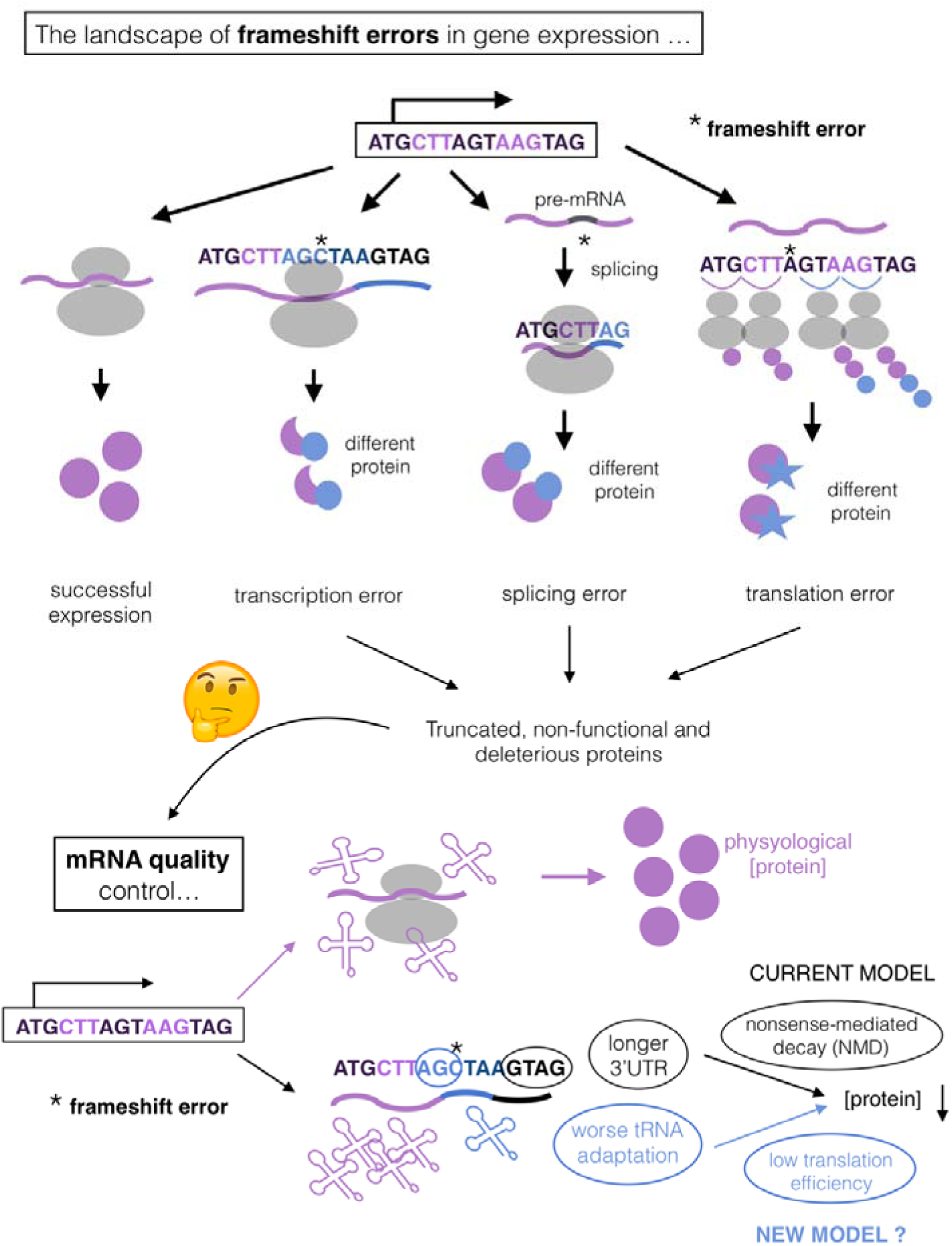
The meaning of codon bias in the transcriptome. **(A)** Highly expressed genes are often selected to have optimized codons in agreement with the cellular tRNA pool, allowing efficient translation of them (purple). This is known as “translational selection” (20–23). On the other hand, genes with a poor codon optimization are inefficiently translated and targeted for mRNA decay (blue) (18). **(B)** Top: in yeast, most native *genes* (purple) exhibit a tRNA Adaptation Index (tAI, as a measure of codon optimality) in the range of ORFs predicted from random transcription throughout the genome (blue). Such random ORFs simulate the absence of codon bias in terms of tRNA adaptation. A small fraction of *genes* have non-random tAI, which corresponds to genes “selected for translation”. Bottom: most native *transcripts* (purple) have high tAI, as compared to random ORFs (blue). This histogram was generated weighting each gene by mRNA expression level (which is exponentially distributed), which indicates the per-transcript distribution of tAI.

### The meaning and role of codon bias

All transcriptomes exhibit imbalances in the synonymous codons used for each amino acid. Not all synonymous codons are equally abundant, a phenomena called “codon bias”(20,21). Highly expressed genes use codons translated by abundant tRNAs (22) and are coded by optimized codons **(Figure 2)**, leading to efficient protein synthesis. Highly expressed genes with efficient translation initiation but with suboptimal codon usage are deleterious and affect the expression of the rest of the proteome (23).

It was previously noted that use of optimal codons increased not only protein levels, but also mRNA levels (24–26), suggesting that ribosome speed might regulate mRNA stability. Recently, a pathway that involves the DEAD-box RNA helicase Dhh1 was found to target transcripts with suboptimal codon usage for decay in a translation-dependent manner (18,27). Even short stretches of twelve suboptimal codons reduce mRNA levels (19), likely due to slower translation (28).

While most genes do not have highly optimized codon usage, the majority of the yeast transcriptome is populated by highly optimized mRNAs (**Figure 2B**). The top 10% of expressed genes have highly optimized codon usage. In yeast these genes account for 77% of the transcripts in a cell. Translational selection (29) will result in the optimized codon usage of constitutively highly expressed genes but will act less efficiently on genes with lower expression, genes that are rarely expressed, and of course on out-of-frame codons.

### Codon optimality for removing frameshifting errors

In addition to producing PTCs, frameshifts are likely to introduce a stretch of non-optimized codons at the 3’end of the ORF **(Figure 1)**. In genes with optimized codons, this will result in a sudden changes in translation efficiency after the frameshift, which will reduce protein synthesis and target the transcript for decay (**Figure 3A**). This reasoning follows the observation that the impact of low codon optimality on translation efficiency and mRNA decay is local and can act over as few as twelve codons (19,28). The magnitude of the decrease in codon optimality will be highest for transcripts with high codon optimization (most of the mRNAs in the cell **(Figure 2B)**), which correspond to highly expressed genes that likely bear most of the frameshifts (assuming a uniform distribution of errors across transcripts (1)). Our hypothesis is that frameshift-removing mechanisms are especially relevant for such highly-expressed genes. Furthermore, the impact of low codon optimality close to the 3’ end of the mRNA is higher (Mishima and Tomari 2016). In the case of a frameshift, the enrichment of non-optimal codons should be towards the end of the ORF, which predicts that the destabilizing effect will be even stronger.

**Figure 3:**
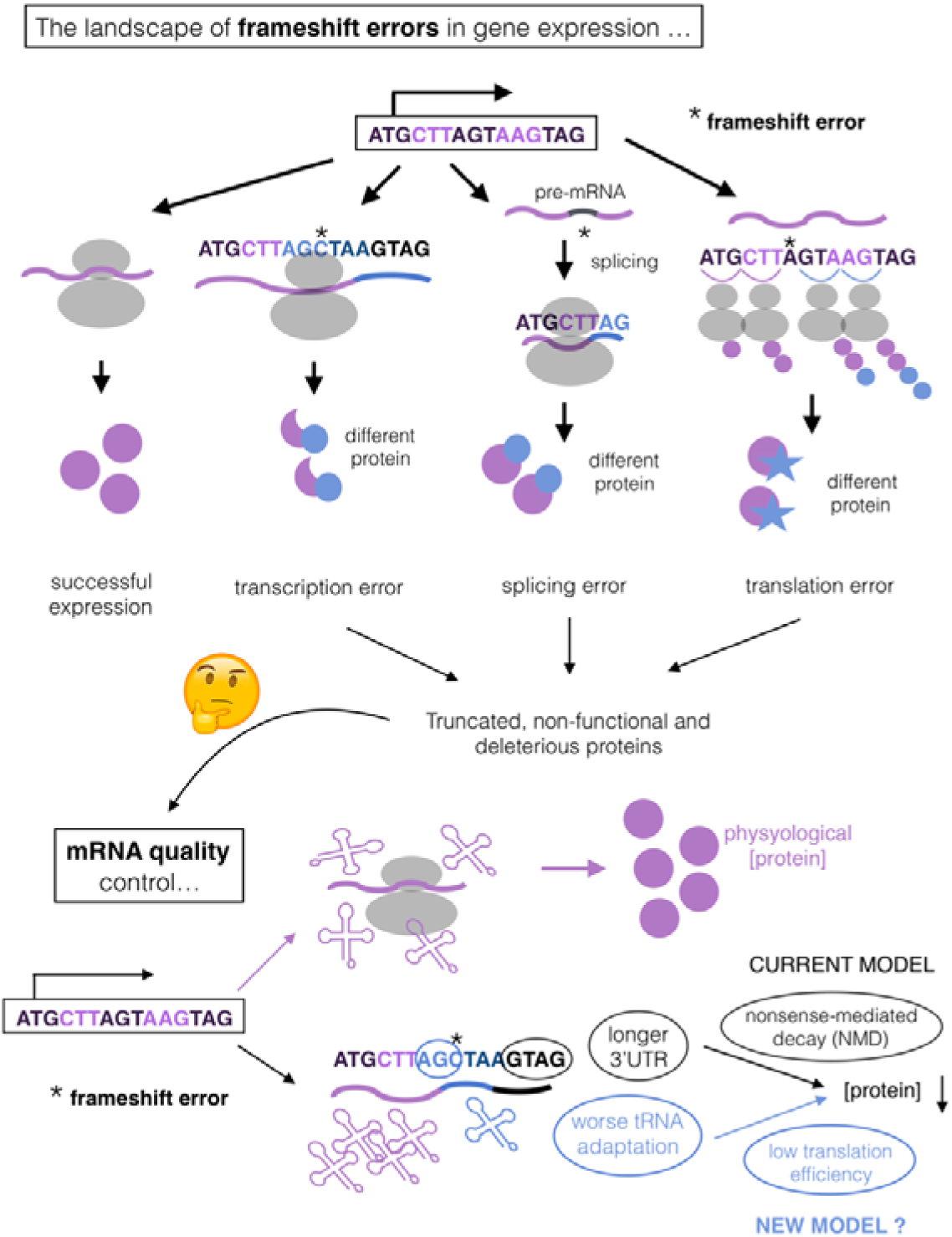
Codon bias can implement quality control of mRNAs with frameshifts. **(A)** tAI follows a negative sigmoidal relationship with mRNA expression levels. Expression was calculated as the log_2_-ratio between mRNA and DNA abundance of a synthetic ORF library of random fragments from the yeast genome, expressed in a plasmid (11). The dashed line represent a threshold in which decreasing tAI reduces expression. **(B)** NMD strength follows a positive linear relationship with 3’UTR length. NMD was measured as the expression (calculated as in A) log_2_-ratio between identical ORF libraries built in a *Δupf1* or a *wt* strain (11). This ratio indicates the impact of NMD for each sequence in the library (which has variable 3’UTR lengths), as UPF1 is responsible for NMD (10). The dashed line represent a threshold in which increasing 3’UTR generates NMD (positive values in the Y axis). **(C)** A pipeline for predicting the impact of NMD and codon on frameshift quality control. As an example of frameshift, we simulated 10^5^ random single-base deletions on native transcripts. Each gene includes a number of mutations proportional to its expression level. For each error (and corresponding native transcript) we calculated tAI between the frameshift and the PTC (local tAI) and the resulting 3’UTR length. We used these as measures of the impact of error on translation efficiency and/or NMD targeting. **(D)** Transcripts with frameshifts (blue) have lower tAI (top) and longer 3’UTRs (bottom), when compared to native mRNAs (purple). The dashed lines represent the thresholds described in A,B.

To compare the role of NMD and codon bias in mRNA quality control we ran a frameshift-introducing simulation on yeast transcripts. We generated random single-base deletions in native transcripts and calculated codon optimality (tRNA adaptation index, tAI (30)) and 3’UTR length with and without the frameshift. Because errors occur on a per transcript basis, each gene received a number of errors proportional to its mRNA expression level (**Figure 3C**).

We found that almost all frameshifts produce a large decrease in tAI after the mutation (**Figure 3D**). The change in tAI range due to frameshifts decreases mRNA levels (11) **(Figure 3A)**. In contrast, ~50% of errors produce 3’UTRs in the range of native 3’UTR lengths (**Figure 3D**), likely unaffected by NMD (11) **(Figure 3B)**. These findings indicate that selection for codon-optimality (which acts on highly expressed genes) can be a robust way to define “correct transcripts” and thus remove transcripts that contain frameshifts

## Discussion

Cells needs to remove transcripts with errors; mutants with increased error rates or that are unable to remove transcripts with errors grow slowly (1,31). Frameshift errors are likely to be deleterious, both by generating deleterious protein isoforms, and because suboptimal codons titrate away both tRNAs and ribosomes (23,32). However, both the sequence features that cells recognize and the mechanisms by which they do so remain poorly understood. Many open questions remain.

NMD is weak (11,12) and affects 5-20% of the native transcriptome (13), so it may be both inefficient and unspecific for removing errors. Removing transcripts with low codon optimality may be more accurate and efficient. This is consistent with the fact that NMD strength follows a linear relationship with 3’UTR length, while codon optimality has a sigmoidal impact on expression **(Figure 3)**. Small changes in codon optimality can lead to a large decrease in expression.

We observe that ~50% of frameshifts generate 3’UTRs within the range of native transcripts, likely unaffected by NMD. This exemplifies how a model based on a qualitative basis (“NMD removes frameshifts *because* these have longer 3’UTRs”) can fail to predict of the quantitative behavior of a system.

Our recent work suggests a genetic link between codon bias and NMD (11). Here we report a possible explanation of this interaction, but it remains to be seen which is the impact on measured expression levels of both processes. The mechanism of this link also remains to be established.

In frameshifted mRNAs, the quantitative impact of the low-tAI stretches of ORF in expression remains elusive. It will be interesting to see if they can explain more or less quality control than NMD. In addition, the effect of codon bias on expression is expected to impact protein levels (20,23), not only mRNA. This predicts that the impact of codon bias on expression is higher than reported here **(Figure 3A)**, which is not true for NMD. This could explain why we observe a lot of splice isoforms that have PTCs in humans, which may arise from frameshifting splicing errors. NMD does not remove them (as we can detect them), but it is likely that they have lower codon adaptation and reduced protein levels.

Finally, this work raises a possible explanation for an adaptive benefit of imbalanced tRNA repertoires (22), which would confer the ability to degrade transcripts that are not supposed to be highly expressed. It is almost certain that cells avoid selecting the expression of ORFs with a random composition of codons. Frameshifts generate such random stretches, that are likely targeted for decay. Thus, there may be an evolutionary pressure for imbalanced tRNA repertoires to ensure proper mechanisms of mRNA quality control. It will be interesting to determine if this process has driven the evolution of codon bias and codon-usage associated mRNA stability, or it is a passive result due to the fact that almost any frameshift will reduce the optimality of the already very optimal genes.

## Methods

### Codon bias measurements

Codon bias was approximated by calculating the tRNA adaptation index (tAI) (dos Reis et al. 2003) for each open reading frame (ORF, either native of the yeast transcriptome or simulated). In order to generate random ORFs we simulated random transcription start sites (TSS) across the whole genome of *Saccharomyces cerevisiae* (Cherry et al. 2012) and generated the ORF starting at the first ATG from the TSS. tAI was calculated on each of them in order to measure the codon bias of random coding sequences.

### mRNA expression weighting

In order to approximate *per-transcript* distributions (of tAI and 3’UTR length) we weighted each gene by the sum of the TPM expression obtained from multiple RNA-seq experiments (generated in (Carey 2015)). This means that each gene has a weight in the distribution which is proportional to it’s mRNA expression.

### Relationship between ORF features and expression

We obtained data about the relationship between several ORF features (3’UTR length and tAI) and mRNA expression from an existing dataset (Espinar et al. 2018). It includes the expression measurements for a library of ~10,000 ORFs randomly generated from the yeast genome. In order to determine the impact of 3’UTR length on NMD we generated generated the same library on a UPF1 deletion strain, as described before (Espinar et al. 2018).

### Simulating frameshifts

As an example of frameshift, we simulated 10^5^ random single-base deletions on yeast native transcripts. Each gene includes a number of mutations proportional to its expression level (as explained in *mRNA expression weighting).* For each error (and corresponding native transcript) we calculated tAI between the frameshift and the PTC (local tAI) and the resulting 3’UTR length.

### Data availability

All code and data are at https://github.com/MikiSchikora/CodonBias_QualityControl

### Funding details

This work was supported by Ministerio de Economía y Competitividad (MINECO) and the Fondo Europeo de Desarrollo Regional (FEDER) BFU2015-68351-P (MINECO / ERDF EU), AGAUR (2014SGR0974 and 2017SGR1054) and the Unidad de Excelencia María de Maeztu, funded by MINECO (MDM-2014-0370).

